# Whole slide image representation in bone marrow cytology

**DOI:** 10.1101/2022.12.06.519318

**Authors:** Youqing Mu, H.R. Tizhoosh, Taher Dehkharghanian, Clinton JV Campbell

**Author notes:** Corresponding author: Clinton JV Campbell.

## Abstract

One of the goals of AI-based computational pathology is to generate compact WSI representations, identifying the essential information required for diagnosis. While such approaches have been applied to histopathology, few applications have been reported in cytology. Bone marrow aspirate cytology is the basis for key clinical decisions in hematology. However, visual inspection of aspirate specimens is a tedious and complex process subject to variation in interpretation, and hematopathology expertise is scarce. The ability to generate a compact representation of an aspirate specimen may form the basis for clinical decision support tools in hematology. We have previously published an end-to-end AI-based system for counting and classifying cells from bone marrow aspirate WSI. Using deep embeddings from this model, we construct bags of individual cell features from each WSI, and apply multiple instance learning to extract vector representations for each WSI. Using these representations in vector search, we achieved 0.58 ± 0.02 mAP@10 in WSI-level image retrieval, which outperforms the Random baseline (0.39 ± 0.1). Using a weighted k-nearest-neighbours (k-NN) model on these slide vectors, we predict five broad diagnostic labels on individual aspirate WSI with a weighted-macro-average F1 score of 0.57 ± 0.03 on the test set of 278 randomly sampled WSIs, which outperforms a classifier using empirical class prior probabilities (0.26 ± 0.02). We present the first example of exploring trainable mechanisms to generate compact, slide-level representations in bone marrow cytology with deep learning. This method has the potential to summarize complex semantic information in WSIs toward improved diagnostics in hematology, and may eventually support AI-assisted computational pathology approaches.

## 1 Introduction

Making a diagnosis is increasingly complex, relying on information from multiple data modalities [1]. Neoplasms are now classified hierarchically based on molecular, clinical and other “multi-omics” data sources [2, 3], but morphology, or the assessment of microscopic features of preserved human, remains the foundation of a diagnosis in pathology [4]. This is based on the fundamental paradigm that semantic information about human disease can be extracted from tissue morphology. However, the way morphology is analyzed is now changing, as the field gradually shifts to using digital whole slide images (WSI) of pathology specimens for morphological assessment. This is called digital pathology, and is poised to become the standard of care for primary diagnosis [5–7]. One proposition of digital pathology is that WSI provide the substrate for computational pathology tools supporting the assessment of tissue morphology [8]. This allows for the extraction of information from tissue specimens, supporting a quantitative and structured approach to morphology, and may yield morphology-based clinical decision support tools (CDST) in pathology workflows. [9].

Artificial intelligence (AI) has been proposed as a method for morphology-based CDST in pathology and other medical disciplines [10]. In its current state, AI consists of numerous techniques, most of which are collectively called machine learning (ML). ML algorithms have the ability to learn inherent patterns in data and generalize these patterns to make predictions on new unseen data, albeit in a narrow domain [11]. The use of AI in analyzing pathology WSI, specifically deep learning, is well-established in terms of histopathology applications such as tumour identification and grading, and tissue biomarker-based prediction and prognosis [12–15]. However, CDST would ideally have applications beyond tumour grading or cell classification, where collective morphological information from a WSI may be extracted, processed, and even interpreted for a pathologist or end-user as a compact representation. This would represent an evolution from an informationextraction-based computational pathology tool to a CDST demonstrating some level of semantic-like tissue interpretation.

In hematology, bone marrow aspirate cytology is essential for clinical decisions with an immediate impact on patient management, e.g., blast counts in acute leukemia [16]. The basic aspirate cytology workflow consists of several steps; first, counting and classifying hundreds of individual cells into a number of categories (called nucleated differential count, or NDC); next, interpreting this count in the context of subtle cytological features (morphology); and finally, interpreting cell counts and features in the context of additional histopathology, molecular, flow cytometric and clinical data to make a final overall interpretation (integrated reporting) [17]. Each of these steps requires years of subspecialized hematopathology training, which is an increasingly limited resource, and there is variation in interpretation by hematopathologists inherent in every step [18–21]. Collectively, these factors may adversely impact patient outcomes, and consequently, clinical decision support tools CDST to support aspirate cytology are highly desirable [10]. Both the NDC, as well as visual semantics from individual cells, form the basis for a predictor of clinical states, e.g., acute leukemias or myelodysplastic neoplasms (MDS) [22]. As a result, the NDC provides a basic but useful description of a slide. To this end, we have recently published an end-to-end AI-based system for counting and categorizing cells from bone marrow aspirate WSI [23]. This allows for the sampling of cells from WSI and generating a probability distribution called *Histogram of Cell Types* (HCT). However, in theory, computational tools that account for the individual cell features, in addition to simple cell counts, would perform better as a CDST, particularly in diagnoses such as MDS, which rely heavily on individual cell morphological features versus cell counts [2, 19, 24, 25]. This *“cell features plus cell counts*” approach would entail a semantic slide-level interpretation that more closely resembles the approach of a pathologist. So far, such approaches have not been implemented in cytology.

Generally, the aspiration to design computational pathology tools that extract conceptual or semantic information from all or part of a WSI to render a slide-level interpretation is called *whole slide representation* [26]. A whole slide representation would ideally be coupled with information from other sources to support clinical decision-making (e.g., clinical, molecular data), a process generally known as multi-modal learning [27–29]. There are similarities in this scheme to how an actual pathologist works: extracting and presenting complex visual semantics from an image, and using this information in the context of other information to construct a “patient representation” to assist the clinical decision-making process. To be clear, the goal of whole slide representation is not to make a final diagnosis, but to allow for a relative comparison of morphological semantics from individual patient WSI. This information can then be used in the context of other information to support the multi-modal diagnostic process.

Several unique challenges have created an impediment to slide-level representation. One of the most significant is the problem of how to represent a large (gigapixel range, image resolutions can be as large as 250000 × 150000 pixels) WSI in a way that efficiently captures the inherent complex visual (and spatial) tissue semantics. Conceptually, again modelling from how an actual pathologist works, visual semantics in smaller image subunits, or patches, would need to be extracted and then pooled to render a representation. This leads to the problem of patch labelling, how individual patches can be labelled or annotated, and which patches should be emphasized in the overall WSI representation. Because slide-level labels may only reflect part of the WSI (e.g., regions containing tumours), most techniques depend on pathologists’ region of interest (ROI) annotations to capture the nuances in different tissue regions [30, 31]. Unfortunately, this procedure is not usually practically feasible, requiring labour-intensive manual labelling, and is also not applicable to a WSI that may have few ROIs.

One possible solution to the problem of patch labelling in WSI is to organise WSI patches in “bags”, with each bag containing a collection of regions with a single label [32]. Different individual labels exist for the instances (patches) within a bag, but collectively they form a single bag label. In basic multiple instance assumption, a bag is regarded as negative if all of its instances are “negative”. On the other hand, a bag is positive if it contains at least one “positive” instance. Individual labels exist but remain unknown during training [33]. As such, detailed individual patch labelling is not required. This is conceptually similar to the procedure of cancer diagnosis: if a cancerous region is identified in a tissue specimen, the diagnosis is cancer, regardless of whether normal tissue is present. Generally, this approach is called multiple instance learning (MIL), a type of weakly supervised deep learning [34]. However, MIL is more data-intensive than training from manually labelled patches [35, 36]. To help mitigate this problem, some studies have applied an attention mechanism with MIL, including attention-based pooling [37, 38], Hopfield attention [39], and selfattention [40, 41]. An attention mechanism is a trainable technique for significantly increasing the efficiency and accuracy of information processing by prioritizing certain data elements and de-prioritizing others [42]. In MIL, an attention mechanism generally assesses all instances of a bag, providing an attention score to each instance that informs its contribution or relevance to the overall embedding representation for a given bag label [42]. Recently, a clustering-constrained-attention multiple-instance learning (CLAM) [43] method used small WSI patches, which have high information density as they represent solid tissue, as inputs to create useful WSI representations in histopathology.

In comparison to histopathology, blood and bone marrow cytology are distinct, where morphological analysis is based on *counting and classifying discrete objects* (*cells of interest*) within patches [23, 44, 45], rather than the whole regions of patches. As a result, many patches in cytology are “useless” (due to the lack of useful cells) or have low information density (the “environment/background” of cells doesn’t provide helpful signal). Under these conditions, in cytology, *any method that uses patches as the basic inputs may fail to train models accurately or efficiently*. One feasible alternative is to use *individual cells* as the base input to extract information from a cytology slide, rather than ROI patches for model training, which are arbitrary crops from WSIs. Using individual cells’ embeddings, MIL bags can be created based on cells with an NDC implicitly included in each bag, and get rid of irrelevant cell background. In this way, the information density of input increases and thereby the challenge of “sparsity” in cytology patches is addressed. Furthermore, cells can be resampled to create new MIL bags with the same count feature, which can serve as a data-augmentation technique to address the issue of insufficient data.

In this work, we present a method to condense cytological information from a bone marrow aspirate WSI into a single, slide-level vector by (a) using embeddings of individual cells as input units instead of WSI patches; (b) resampling cells from a WSI to construct MIL bags while preserving the NDC; (c) applying attention-pooling on the cell bags (Fig. 1). Our results indicate this approach can work well with minimal data and is robust against noise, and is computationally efficient. It generates a compact WSI representation that may, in principle be used to support simple, slide-level diagnosis, or patient comparison, in hematology. To our knowledge, this proof-of-concept work is the first study applying MIL in cytology, and using individual cells as input to create a compact WSI representation in pathology, which may eventually form the basis for a CDST in hematology when considered in the context of other data sources. Furthermore, our method is interpretable, where class-associated cells can be extracted and visualized to gain biological insight and understand the model’s working mechanism.

**Figure 1:**
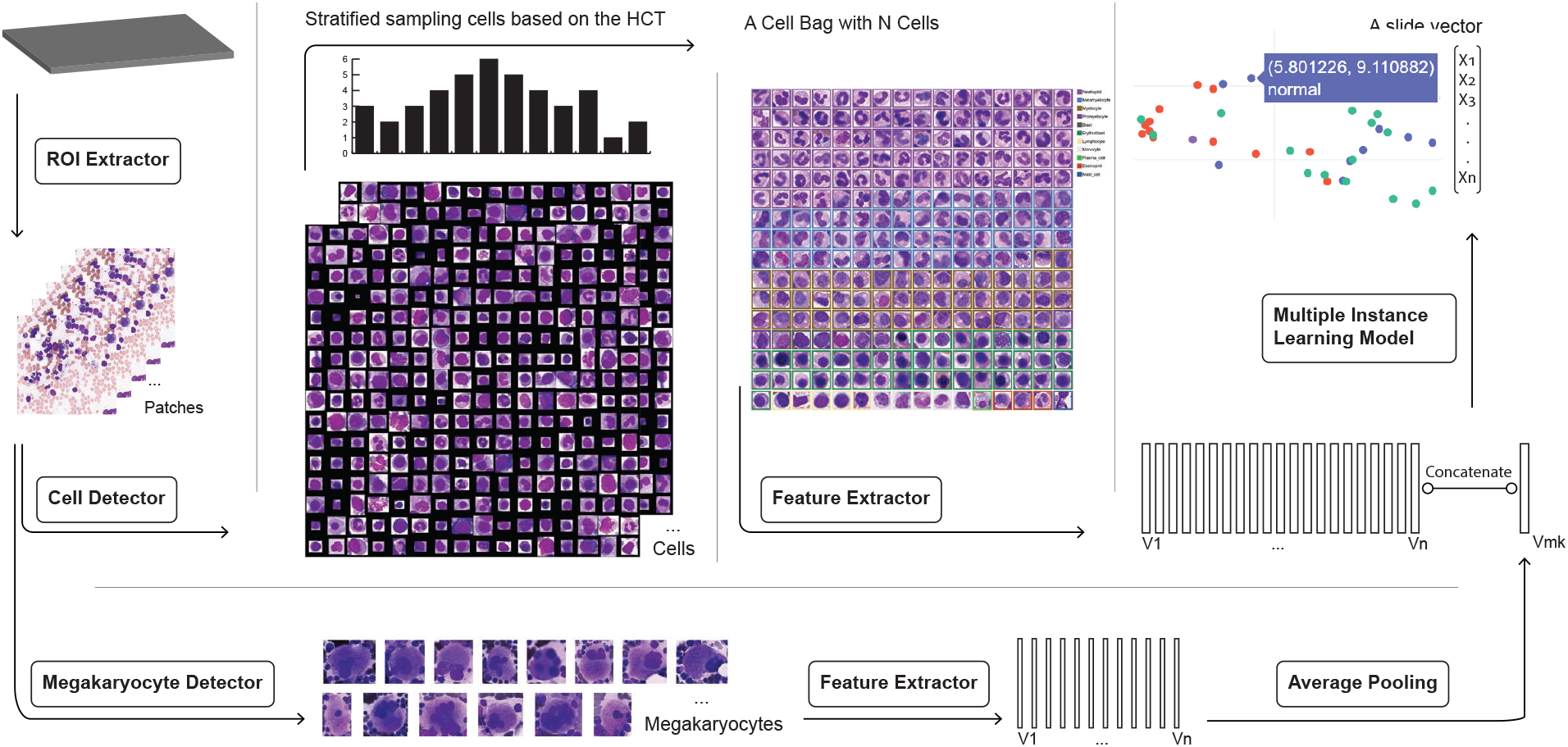
The end-to-end process. Using our previously published system for aspirate cytology, regions of interest (ROIs) from each WSI were identified, and then individual cells are detected in each ROI. Subsequently, cell bags for each WSI were created from individual cells (bags can be resampled multiple times). The hidden layers from a fine-tuned YOLO model were used as a feature extractor for individual cells. Megakaryocytes went through a similar process, with several modifications due to their uniqueness (see methods). Finally, these cell features were concatenated with the megakaryocytes’ vector and input through a model that integrates them into a single slide vector.

## 2 Methods

### 2.1 Dataset

WSIs were acquired retrospectively de-identified and annotated with only a synopsis, spanning a period of 1-year and 865 patients. Glass slides were scanned at 40X magnification. The data comprised a full range of diagnoses at a large hematology reference centre throughout this time period. The study was conducted under Hamilton Integrated Research Ethics Board (HIREB) study protocol 7766-C.

We previously deployed an end-to-end YOLO-based technique for detecting, classifying and counting cells from bone marrow aspirate WSIs [23]. 556 WSIs which have more than 256 cells that are in the label set of {Neutrophil, Metamyelocyte, Myelocyte, Promyelocyte, Blast, Erythroblast, Megakaryocyte nucleus, Lymphocyte, Monocyte, Plasma cell, Eosinophil, Basophil, Histiocyte, Mast cell } were selected.

Labels were created by simplifying the predictions of a fine-tuned BERT model on WSI synopses, as previously reported by our group [46]. This BERT model’s predictions took the form of a multi-label task. In this work, these predictions were simplified to broad diagnostic categories using the following rules:

1. If “normal” is in a multi-label prediction, this prediction will be simplified as “normal”, e.g. “normal; iron deficiency” becomes “normal”.
2. If any label is “acute leukemia” related, this prediction will be simplified as “acute leukemia”, e.g. “acute myeloid leukemia; acute promyelocytic leukemia; hypercellular” becomes “acute leukemia”
3. If any label is “myelodysplastic syndrome” related, this prediction will be simplified as “myelodysplastic syndrome”, e.g. “hypercellular; myelodysplastic syndrome” becomes “myelodysplastic syndrome”.
4. If any label is “plasma cell neoplasm” related, this prediction will be simplified as “plasma cell neoplasm”, e.g. “hypercellular; plasma cell neoplasm” becomes “plasma cell neoplasm”.
5. If any label is “lymphoproliferative disorder” related, this prediction will be simplified as “lymphoproliferative disorder”, e.g. “hypercellular; lymphoproliferative disorder; mastocytosis” becomes “lymphoproliferative disorder”.

Predictions that did not fit the aforementioned conditions have less than 10 cases in their groups, which is insufficient to form new classes. We therefore discarded 61 such cases. In this way, we created a primary dataset of 556 WSI and label pairings.

### 2.2 Stratified sampling cells using rHCT as cell bags

This aggregate cell count information from the cell detection system was collected in a representation called Histogram of Cell Types (HCT), which quantifies the probability distribution of bone marrow cell classes [23]. We limited its included cell types to {Neutrophil, Metamyelocyte, Myelocyte, Promyelocyte, Blast, Erythroblast, Megakaryocyte nucleus, Lymphocyte, Monocyte, Plasma cell, Eosinophil, Basophil, Histiocyte, Mast cell } only, called *rHCT* (Refined HCT)). We then used each WSI’s rHCT to guide the stratified sampling [47] process. This allowed us to gather a small number (256) of cells that included the cytomorphological information and cell type probability distribution of a WSI (rHCT), called a “cell bag”.

### 2.3 Re-purposing YOLO as cells’ feature extractor

To extract cell characteristics from cell bags, we used the outputs from the YOLO’s feature extractor layers. Because the YOLO model was trained and validated [23] on bone marrow cytology samples, its feature layers are reliable to capture the salient morphological features of the detected cells.

One patch image (608 × 608) was first transformed into a (76, 76, 256) array via YOLO feature extractor layers [48]. Then we located the bounding box of each cell on this patch’s array to retrieve the array representing that cell. For example, a cell with a bounding box of (x1 = 8, y1=8, x2 = 80, y2 = 80) has its feature array on this patch’s array with a bounding box of (x1 = 1, y1 = 1, x2 = 10, y2 = 10) as the tile image resized smaller for a factor of 8 under YOLO feature layer. The cell array was then averaged over its first and second dimensions to generate its final feature. In the above example, the array (9, 9, 256) was averaged over its first and second dimensions to produce a (256,) vector representing a single cell. This process resulted in a cell bag (a matrix with the shape of (256, 256)) that consists of cell features.

### 2.4 Concatenation of features from megakaryocytes with cell bags

Megakaryocyte detection was addressed differently in terms of model design, since they are significantly bigger and rarer than other cells. As a result, we employed a different object detection model than YOLO to detect megakaryocytes throughout WSI. Because megakaryocytes are uncommon in comparison to other cell types, the entire WSI was examined in order to locate a sufficient quantity of megakaryocytes. Thus, an ROI detection model was unnecessary. To this end, we trained another model (CenterNet HourGlass-104 512 × 512) [49] on a lower magnification as an object detector to find megakaryocytes more efficiently. This model’s window size was 16 times larger than that of YOLO, with an average precision of 0.86. The YOLO feature extractor was then utilised to compute megakaryocyte features. We averaged their features in a WSI and obtained a vector of size (256,) which was then concatenated with the features of a cell bag. Their concatenation produced arrays of size (257, 256), which served as the *input* for the MIL model(Fig. 1 2).

**Figure 2:**
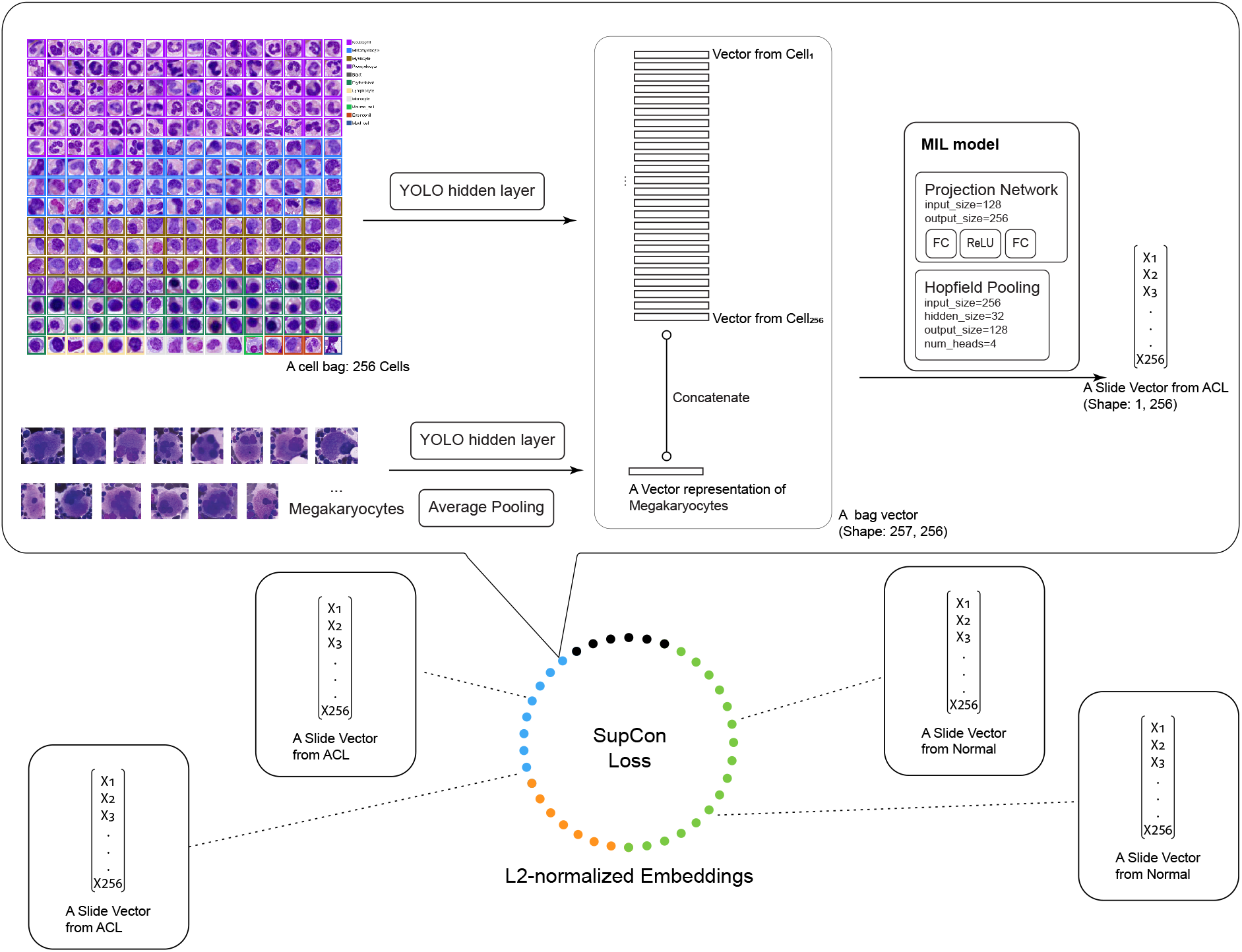
The modelling process. Supervised contrastive learning (SupCon) was used to direct the model training process. Its loss function “pulls together” a case and the cases with the same label in the embedding space, and “pushes apart” the case from cases with different labels. Using this method, the multiple instance learning model learned to generate slide representations that are diagnostically meaningful from cell bags.

### 2.5 MIL with modern Hopfield networks and metric learning

Modern Hopfield networks act as transformer-like attention mechanisms, which in contrast to classical Hopfield networks, are generalized to continuous states [39, 50]. The modern Hopfield network’s update rule is known as the key-value attention mechanism (Equation 1), which has proven to be particularly successful in natural language processing via the transformer and BERT model [51, 52]. Furthermore, modern Hopfield networks have exponential storage capacity in the feature space dimension and converge after just one update, which allows them to extract patterns among a large set of instances [39]. For our study, the Hopfield attention mechanism allowed the network to learn clearly for each label which cell’ morphological features should be regarded as positive evidence (predictors of the slide label) vs. negative evidence (non-informative) and summarise distinct slide-level representations.

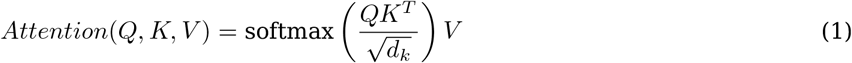

In further detail, a Hopfield-pooling with four-head attention (Equation 2) and a small projection network with two fully connected layers were used in our pipeline to produce a slide-level representation for each WSI.

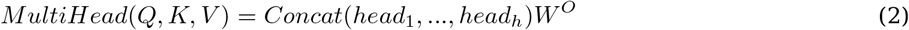

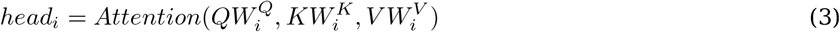

We trained this model via metric learning using PyTorch Metric Learning [53] library’s MetricLossOnly trainer with early-stopping on training loss (patience=5). The goal of metric learning is to generate an embedding space in which related objects are close together while dissimilar ones sit far apart [54]. A common strategy is to use the triplet [55, 56] or N-pair losses [57, 58]; the former employs only one positive and one negative sample per anchor, while the latter uses one positive and many negatives. But both need hard negative mining, which can be challenging to tune well. To avoid the necessity for hard negative mining, our model’s weights were trained with supervised contrastive (SupCon) loss 4, a supervised learning loss that uses many positives and many negatives for each anchor and thus eliminates the need for hard negative mining[59].

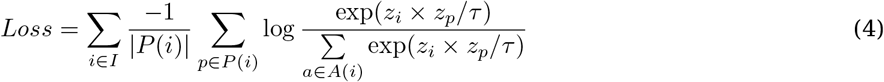

The structure of this approach is similar to that of self-supervised contrastive learning, with adjustments for supervised categorization. Instances go through an encoder network, and the resulting embeddings are then L2-normalized and go through a projection network (Fig. 2). The projection network’s normalised outputs are used to compute the SupCon loss. As such, the normalized embeddings from the same class are drawn together more closely than embeddings from different classes[59]. SupCon loss method outperforms crossentropy method [60] on CIFAR-10, CIFAR-100 and ImageNet. SupCon loss is also robust to noisy labels and more stable to hyperparameter settings like optimizers and data augmentations[59]. Because our labels come from our previously published BERT model’s predictions[46] rather than experts, we must take its prediction errors into consideration. SupCon loss’s tolerance to noisy labels became an obvious option for this training. As such, the model can learn how to generate a vector for a WSI that maximises its “representability” even with noisy labels.

### 2.6 WSI retrieval via vector search

The process of generating a WSI-level embedding can be regarded as a transformation process that derives numeric representation from high-dimension and high-noise sources. The WSI and its embedding form a 1-to-1 mapping relationship. Using approximate nearing neighbor algorithms on these embeddings, semantically close instances can be retrieved, which is called vector search [61].

To retrieve WSI, the distances between an input query WSI and all WSI in the database are calculated and ranked from low to high. Then, the top-k results are directly taken as the returns. mAP@10 (mean average precision at 10) [62] is used as the evaluation metric for our WSI retrieval task.

For a single query:

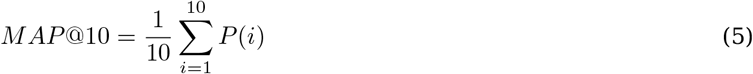

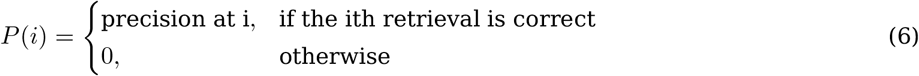

### 2.7 Making predictions on slide vectors by *k*-Nearest Neighbor (*k*-NN)

In our pipeline, to predict the WSI label from slide vectors, we used a weighted *k*-NN classifier with *k*=17 (heuristic 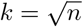, where n is the sample size, 278) and cosine distance

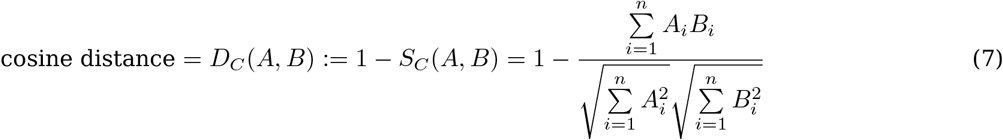

as the distance measure. k-NN can be described as a classification method whose output is a class membership. An instance is categorised by a majority vote of its neighbours, with the instance being allocated to the class most prevalent among its k nearest neighbours. Weighted-*k*-NN [63] discriminatory assigns distinct weights to each of a query point’s k closest neighbours by the inverse of their distance weight 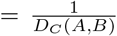. In this case, neighbours who are closer to a query point will have a bigger effect than neighbours who are further away. In contrast to model-based algorithms, this approach is instance-based learning that does not learn weights from training data to predict the output. As a result, it’s a widely used approach for assessing the quality of embeddings [62]. Furthermore, because its inference method is analogous to throwing a dart and the dart’s class is determined by where it falls on the board, the prediction result is simple to understand.

As no published appropriate benchmarks were found, our method’s basic benchmark was a classifier that makes predictions based on a multinomial distribution parametrized by the training class prior probabilities. We used *k*-NN and the empirical classifiers that are implemented in Scikit-learn package [64].

### 2.8 Attention weight plot from Hopfield Pooling

The attention network computes attention weights for cells in a cell bag, which directly represents the importance of each cell [39]. We took the mean of these four head attention weights (Equation 8) as the attention weight of this cell since we employed four heads (Fig. 2) in the Hopfield Pooling, with each head computing its attention weight for each cell feature input.

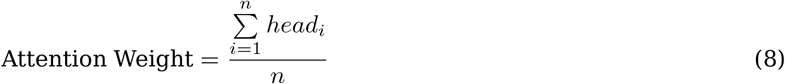

This importance value is an interpretable quantity that can aid in the comprehension of the model and potentially in finding new biological discoveries.

To graphically represent the attention weights for cells in a cell bag, we grouped cells by cell type and ordered these groups by the attention weight sum of each group. Because this ranking is affected by the number of cells in a group, we also provide the average attention weight in a group. However, because the slide-level representations are summarised with all morphological aspects examined using attention weights (Section 2.5), the average attention weight is less important than the attention weight sum. Thus, we ranked the cells by their attention weight sum instead of their attention weight average.

## 3 Results

### 3.1 Visualizing slide representations in the test dataset

We first sought to visualize the quality of the representations generated from the MIL model. Slide representations were generated by the MIL model from features in cell bags. To reduce the influence of unrepresentative sampling during bag creation, we resampled bags 16 times during inference and used the average of 16 bags’ outputs as the final slide representation. We randomly split 556 cytology slides into a training set (50% of cases, i.e. 278) and a test set (50% of cases, i.e., 278) with class stratification and used UMAP to show the slide representations from the test dataset in two dimensions to obtain insight into the diagnostic significance of these slide representations. The key reason for a “0.5-0.5 train-test split” was that our dataset is relatively small, yet a certain absolute number of examples were required to better visualize and comprehend the slide representations. This split method gave enough visualization examples and made the test set results more reliable.

We found that generally, the representations tended to form diagnostically meaningful associations (Fig. 3). For instance, because the “normal” label reflects a broad and heterogeneous patient population, it is not surprising that the representations from these WSI are grouped loosely, but, at the same time, do not overlap clusters of clearly defined disease classes, such as “acute leukemia (ACL)”, “plasma cell neoplasm (PCN)” and “lymphoproliferative disorder (LPD)”. The representations from WSI labelled with binary-like haematological disease states, such as “PCN” or “ACL” cluster quite tightly, as predicted. Cases labelled “MDS” tended to form more heterogeneous clustering, with a subset closer to acute leukemia as expected, given that there is overlap between some MDS subtypes with high blasts and AML. This may reflect the biological heterogeneity and diagnostic process in MDS, which always requires ancillary clinical and molecular data (i.e., multi-modal approach). Together, these results suggested that the WSI representations were diagnostically relevant.

**Figure 3:**
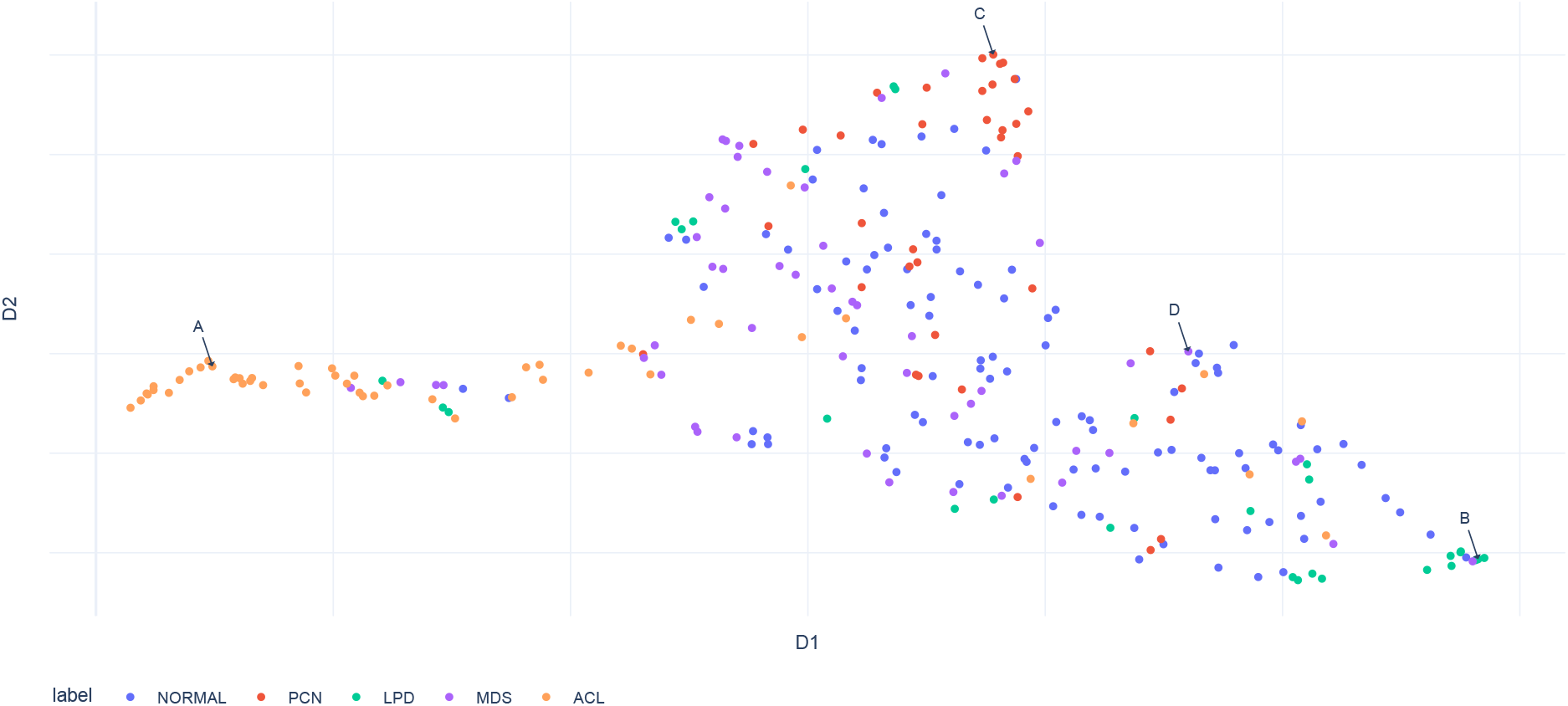
WSI representations, i.e. slide vectors. 2D projection of WSI representations from the 278 cases in the test set is shown. They are colored according to simplified labels. The cases with the same label tended to cluster together, which suggests that the representations are diagnostically meaningful. The special relationship among these representations also captured the natural transition among these disease states. For example, the representations from NORMAL cases locate around the centre and spread among other labels, but they have a clear boundary with ACL. [Readers can also interact with this graph on https://c-campbell-lab.github.io/slide-vector-plot/).]

### 3.2 WSI retrieval using WSI representation

To further assess our model’s capacity to create diagnostically relevant semantic representations, we used 5-fold Monte-Carlo cross-validation to assess the quality of the representations as WSI retrieval problems [65]. For each cross-validated fold, we randomly partitioned 556 cytology slides into a training set (50% of cases, i.e., 278) and test set (50% of cases, i.e., 278) and maintained the proportions of classes consistent in each set.

To enable the WSI retrieval via vector search (Section 2.6), we used cosine distance as the distance measure meant and calculated the distance between the input query (every WSI in the test set) and all WSI in the database (train set). After ranking the distance from low to high, top-ten WSI are retrieved as the query result and used to capture the retrieval quality (Equation 5). As seen in (Fig. 4A), our method achieves 0.58 ± 0.02 mAP@10 in WSI-level image retrieval, greatly outperforming the random retrieval baseline (0.39 ± 0.1).

**Figure 4:**
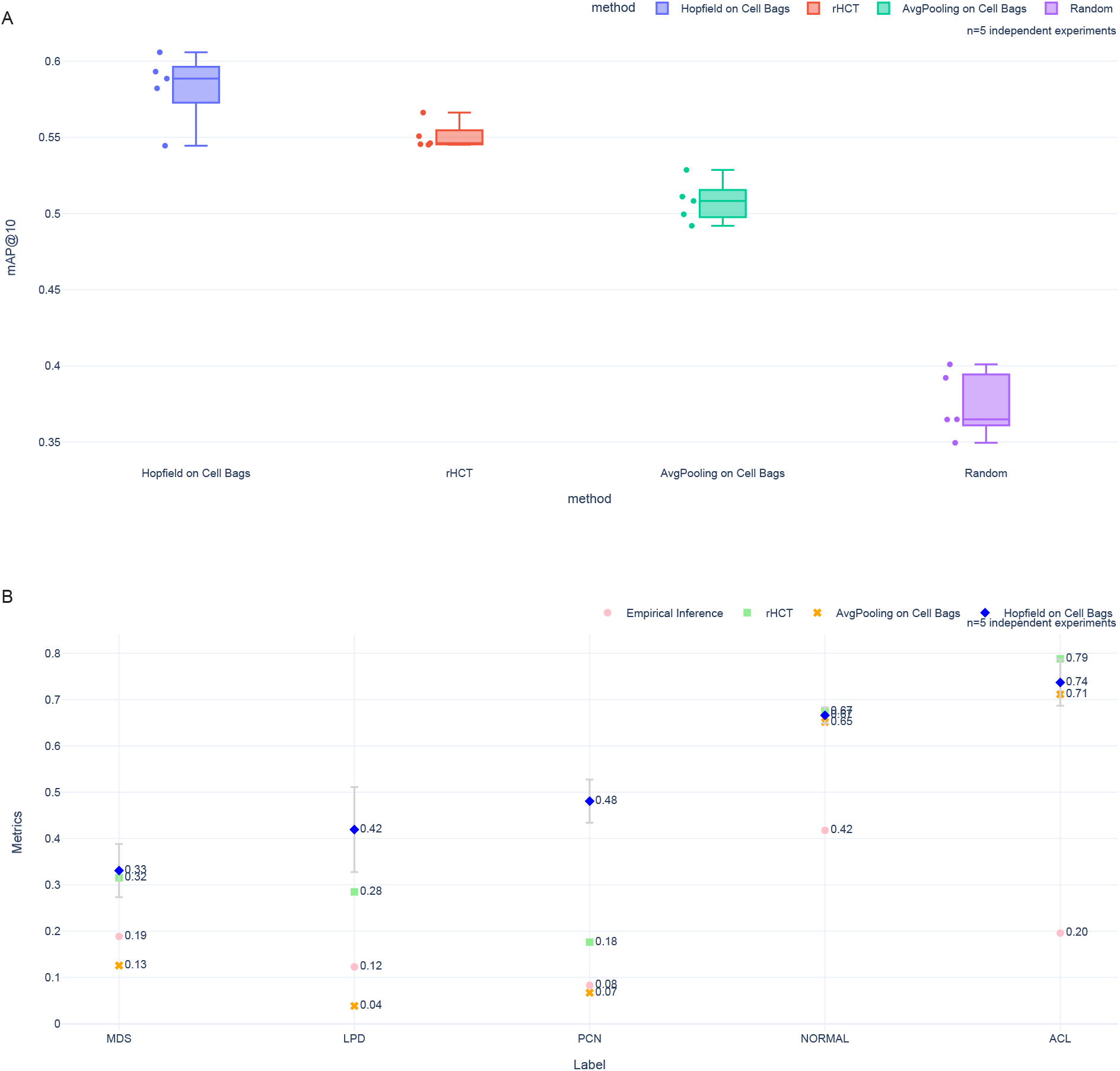
WSI representations quality. A: The mAP@10 of WSI retrieval via vector search on four different methods. This metric measured the WSI representations quality via “search-like” experiments. Only in one experiment did Hopfield on Cell Bags yield an inferior result than rHCT. In all the other experiments, Hopfield on Cell Bags had superior performance than other methods. B: The mean F1 scores and the standard deviation computed across five validation experiments for each simple diagnostic label are shown (Supplementary Table S2). As no published appropriate benchmarks were found, an empirical probability classifier and a k-NN classifier on rHCT were used as comparisons. Our method outperformed the empirical probability classifier and the average-pooling-based MIL algorithm (see Supplementary Figure S1 for the representations from average-pooling). Interestingly, our approach outperformed the k-NN classifier for MDS, LPD, and PCN, suggesting that cell count is a potent predictor of NORMAL and ACL. The F1 scores of each label’s prediction generally reflected the difficulty of diagnosis. For example, ACL has the highest F1 score. It coincides with the relatively simple almost binary diagnosis based on cell counts. Meanwhile, MDS poses a challenge to even experienced hematopathologists and requires multi-model data, and our model struggled with its prediction as well.

To verify whether the attention mechanism improved performance, we switched our system’s pooling component from Hopfield pooling to average pooling and compared the performances before and after this replacement. For this WSI retrieval task, the Hopfield pooling system performed better than the average pooling system in all five experiments (overall, 0.58 ± 0.02 vs. 0.51 ± 0.01, Fig. 4A, Supplementary Table S1). Since average pooling cannot be trained but Hopfield pooling can, replacing the pooling component may be seen as an experiment to assess whether the training process supports better system performance. The result suggested training process indeed improved the WSI representation quality.

To evaluate the improvement brought about by cells’ deep features versus cell counts alone, we also utilised the same WSI retrieval method (Section 7) on rHCT. The rHCT accounts for only the simple counts of cells in each category, and does not account for features in each cell [23], which suggests it should be inferior to the representations from the Hopfield pooling system. As expected, the Hopfield pooling system outperformed this rHCT-based method in four out of five experiments (overall, 0.58 ± 0.02 vs. 0.55 ± 0.01, Fig. 4A, Supplementary Table S1).

### 3.3 WSI classification using slide representation

Under the same experimental setting as the WSI retrieval, we used weighted-k-NN as the classifier to predict the label from slide vectors. This classifier predicted slide-level diagnostic labels with an F1 score of 0.74 ± 0.05, 0.67 ± 0.02, 0.48 ± 0.05, 0.42 ± 0.09, 0.33 ± 0.06 on the test set of 278 randomly sampled WSIs for “ACL”, “NORMAL”, “PCN”, “LPD” and “MDS”, respectively (weighted-macro-average F1 score of 0.57 ± 0.03). Overall, this was superior to the performance of a baseline classifier using a multinomial distribution parametrized by the empirical class prior probabilities with an F1 score of 0.19 ± 0.06, 0.41 ± 0.05, 0.08 ± 0.05, 0.12 ± 0.02, 0.19 ± 0.03 for the same labels, respectively (Fig. 4B). This result suggested that our model efficiently generated simple, diagnostically relevant slide-level representations from bone marrow aspirate WSI.

When verifying the influence of the attention mechanism in the classification task, the Hopfield pooling system performed marginally better than the average pooling system on ACL and NORMAL, but it outperforms on PCN, LPD, and MDS (Fig. 4B, Supplementary Table S2). Again, average pooling cannot be trained but Hopfield pooling can. The Hopfield pooling system’s improved performance demonstrated that training indeed improved performance. Because the parameters in the attention mechanism are trainable via the training data, if more data is available, it has the potential to learn more complex *patterns* in cell bags.

Compared with the rHCT method, our system outperformed this rHCT-based classifier only for MDS, LPD, and PCN, suggesting that cell count is a strong predictor of NORMAL and ACL, while MDS, LPD, and PCN need deep cell features as prediction factors (rHCT in Fig. 4B). This finding is in line with clinical practice, where diagnoses of MDS, LPD and PCN rely more on cell morphology and ancillary (multimodal) data than absolute cell counts, whereas in cases of acute leukemia and normal bone marrows, cells counts alone are highly influential for the diagnosis [66, 67].

### 3.4 Attention on cell types in cell-bags

Each input object, i.e., cell bag, is composed of 256 cell feature vectors and one megakaryocyte average feature (Section 2.4), which are reduced by the Hopfield attention-pooling function to a single 256-length vector representing the WSI. During the process, the attention mechanism within calculated the relevance of these 257 instances in bags as a 257-length “attention weight” vector. This attention value is an interpretable number that indicates the importance of each instance in the bag.

To gain insight into the attention mechanism, we presented the attention weights of the 256 cells in the form of an attention plot (Section 2.8). The last value in the attention weight vector is not used because it is the megakaryocyte average feature, which is not derived from a single cell.

Slide A, B, C, and D (label: ACL, LPD, PCN and MDS, respectively) are interesting cases to examine in detail. These are typical samples located in clusters of acute leukaemia, lymphoproliferative disorder, plasma neoplasm and myelodysplastic syndrome. (Fig. 3). From their attention plot (Fig. 5), we can see in acute leukemia, blast cells receive the highest attention score, as expected (Fig. 5A). Similarly, in LPD, lymphocytes received the highest attention score (Fig. 5B. In PCN, plasma cells received a high attention score, but interestingly, neutrophils received the highest score (Fig. 5C). This may suggest that there are features not currently understood that are present in neutrophils in this case, or, that the model learns to associate increased reactive neutrophils with PCN, which is often the case clinically [68]. In MDS, neutrophils and erythroblasts were weighted the highest, which conceptually makes sense, as the granulocytic and erythroid lineages are critical for the assessment of dysplasia in the diagnosis of MDS [2]. Overall, this suggested that the attention mechanism is interpretable, has diagnostic relevance, and also, that the WSI representation accounted for additional semantic information from cell morphology beyond the simple NDC to make a label prediction.

**Figure 5:**
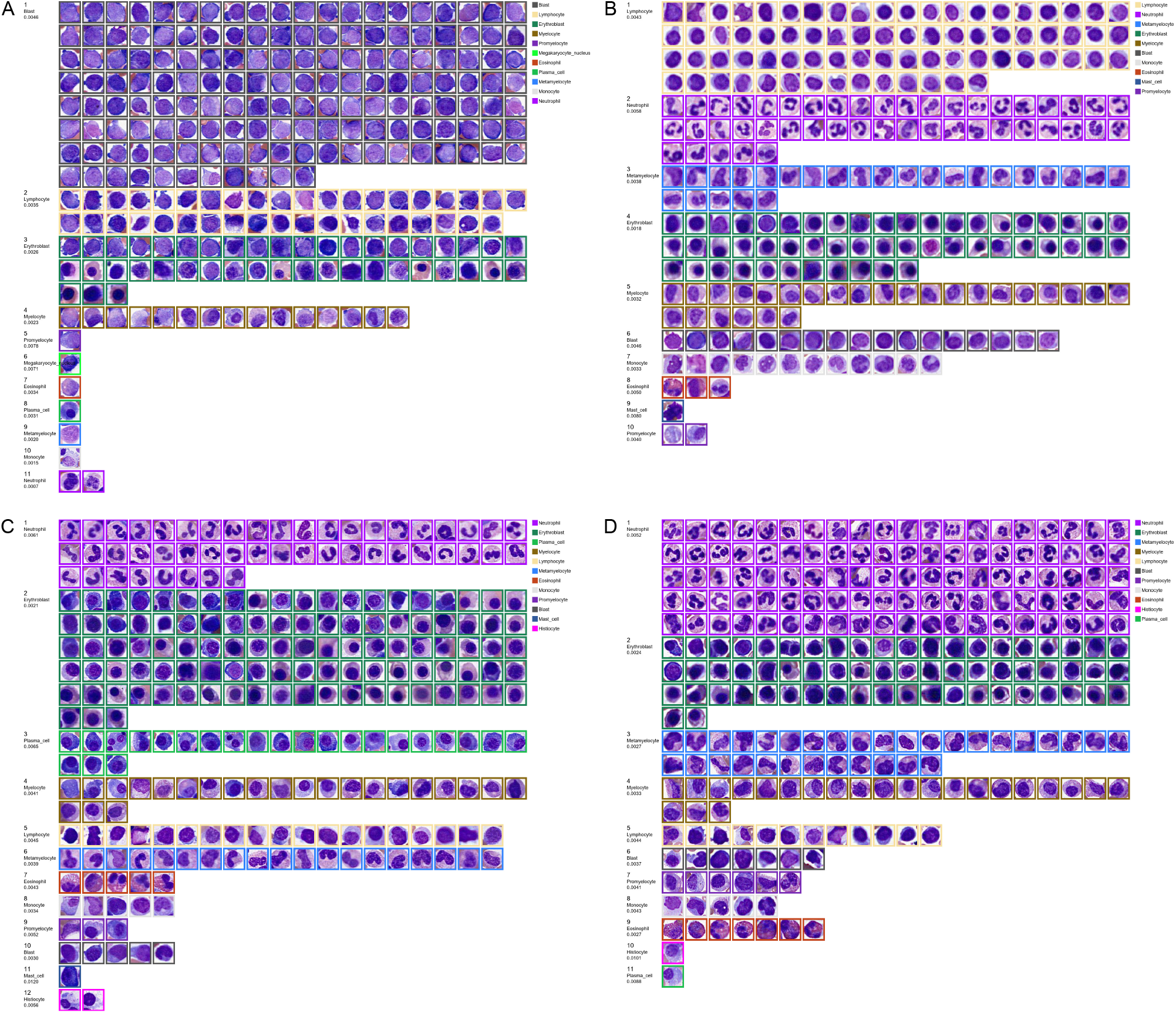
The Attention plot of four cell bags. Attention plots attempt to explain the possible underlying model reasoning process. A, B, C, D have predictions: ACL, LPD, PCN and MDS, respectively. A: blast cells have the highest attention score, as expected for ACL. B: in LPD, lymphocytes have the highest attention score as expected. C: in PCN, plasma cells have a high attention score as expected, but interestingly, neutrophils have the highest score. D: in MDS, neutrophils and erythroblasts are weighted the highest, which conceptually makes sense. Overall, this suggested that the attention mechanism has interpretable. [Readers can view all attention plots by clicking the dots on this interactive graph https://c-campbell-lab.github.io/slide-vector-plot/]

## 4 Discussion

In this paper, we present a computational system that captures diagnostically relevant information as slide-level embeddings from individual aspirate cytology WSI. The novelty of our method is that we created bags using individual cells as the basic inputs rather than patches. This “cell bag” method keeps both the cytomorphological and cell type probability distribution information, while being readily visualized. Our system can generate one slide-level representation from one bag. This representation contains the information of cells from fifteen cell types (“Neutrophil”, “Metamyelocyte”, “Myelocyte”, “Promyelocyte”, “Blast”, “Erythroblast”, “Megakaryocyte nucleus”, “Lymphocyte”, “Monocyte”, “Plasma cell”, “Eosinophil”, “Basophil”, “Histiocyte”, “Mast cell”, “Megakaryocyte”). The information in cell bags lays the groundwork for developing predictors of broad diagnostic labels, and may form the basis for an eventual CDST in hematology.

The most challenging part of our process is for the model to pick up the most relevant “signals” in a small data set. Further, because of the data size limitations, the model may have overfitted the training samples. To help address this issue, we applied several techniques. First, we repurposed our previously reported YOLO model [23] as a frozen feature extractor and turned all cell images into sets of low-dimensional feature embeddings prior to training the model. As the feature extractor (YOLO model) has been tested and can generate high-quality cell embeddings, the total “noise-to-signal” ratio was reduced by using cells’ embeddings rather than raw cell images directly. Second, to reduce the influence of unrepresentative sampling during bag creation, we resampled cell bags constantly throughout model training, and also resampled bags sixteen times during inference. The ability to resample cells to generate new cell bags accomplishes comparable functionality to “Dropout layer”[69] and “data augmentation”[70], which not only helps regularise our MIL model but also improves the information utilisation during training and inference.

In terms of the weaknesses of our system, WSI labels came from our previously published BERT model’s predictions [46] rather than experts, which are not perfectly correct. This was implemented as it is not practically feasible to manually label many hundreds of WSI with keywords from semi-structured diagnostic synopses. Additionally, our YOLO has an error rate during cell detection as described, particularly for blasts and lymphocytes [23]. The errors from the BERT model and the YOLO model will therefore *cascade* through our pipeline, reducing its performance. To address this issue, we trained our MIL model using supervised contrastive (SupCon) loss (Section 2.5) as the loss function. According to the study [59], SupCon loss is more resilient to noisy labels and more stable to hyperparameter settings like optimizers and data augmentations[59]. Meanwhile, our results indicated that our method is resistant to data noise. For WSI retrieval, Hopfield on Cell Bags had better performance than all other methods in almost all experiments (Fig. 4A, Supplementary Table S1). Also, when we utilised weighted-k-NN as the classifier to predict the label from slide representations, the weighted-k-NN outperformed the baseline classifier (Section 3.3) in all labels. These results suggested that we successfully generated high-quality slide-level representations from noisy data. Further work with larger, better-quality datasets will also help to mitigate these issues. Specifically, custom scanning hardware for bone marrow cytology will be required.

In terms of model interpretability, the attention network could compute attention weights for cells in a cell bag, explicitly displaying each cell’s importance. The attention plots were created by combining attention weights with cell bags. When trying to make sense of the system’s prediction, pathologists, for example, could use an attention plot to quickly identify which cells are used by the system. As seen in Figure 5, it is quite simple to determine if the system’s prediction is likely to be correct by using this attention plot. The interpretability feature would be essential for the development of a relevant CDST. In some cases, there were unexpected relationships between cell types given the highest attention, and label prediction by our model (Fig 5). For example, the highest attention in the prediction of PCN was given to neutrophils. This is not totally surprising, and the neutrophil count may be increased in PCN due to reactive change [68]. This may have represented the model learning a relationship between neutrophils and PCN, or alternatively, there may be previously unappreciated subtle features present in neutrophils that were identified by our model, but not readily identified by hematopathologists. This information could help pathologists and researchers readily identify and address the system’s potential problems for further improvement. It may also one day support new biological discoveries.

The performance of the k-NN classifier on rHCT predictions was comparable to that of this deep learning system in some instances. The k-NN classifier on rHCT performed as well as this deep learning system on ACL and NORMAL predictions. This is not surprising, and these diagnostic categories may generally be inferred by hematopathologists by the NDC alone. However, in other diagnostic categories, such as PCN, LPD and MDS, that rely more on cellular features and ancillary clinical and molecular multimodal data versus counts, our deep learning system showed a clear advantage (Fig 3B). We anticipate that with increased WSI quality and quantity, our pipeline will be able to provide better slide representations. Lastly, deep learning systems are renowned for their capacity to enhance as their training dataset expands [71]. Our system was only trained on 278 cases. Its performance is expected to improve as digital pathology becomes more widely adopted and larger, better-quality datasets become available to the community.

In terms of applications, the ability to extract slide-level representations from cytology slides opens the door to many AI-assisted computational pathology applications and CDST. The slide representations, for example, could be utilised to power inter-slide search via vector search, as our preliminary WSI retrieval experiments demonstrated (Section 3.2). Also, as seen from the WSI representations (Fig. 3), complicated cases are more likely to be found along the borders of each cluster. This may be used as a guide for “diagnosis triage,” allowing hematopathologists to devote more time to solving complex cases. Our approach also enables human-centred AI [72], which uses AI as CDST to support the hematopathology diagnostic workflow. Furthermore, slide-level representations can serve as the foundation for WSI-oriented multi-modal analysis. For example, by employing this system as an “image encoder” for WSI, it is feasible to build a CLIP-like system that integrates a huge number of pathology reports or synopses directly with WSI and efficiently learns visual ideas from natural language supervision [73]. Moreover, our technique may be applicable to additional cytology areas (e.g., blood films and body fluids). To this end, we propose an easy-to-understand proof-of-concept technique to describe the system to experts and non-experts alike, by using the slide representation to power vector search and combining the attention plots (Fig 5) with the slide projection plot (Fig 3) similar to an interactive graph https://c-campbell-lab.github.io/slide-vector-plot/. For example, a slide is predicted as ACL since its similar cases retrieved by the search engine are ACL. The attention plot may then be used to determine whether the slide embedding placement of this slide makes clinical sense (Fig 6). In this way, our system may form the basis for an eventual CDST in hematology, which is relevant to diagnosis and ultimately better patient outcomes.

**Figure 6:**
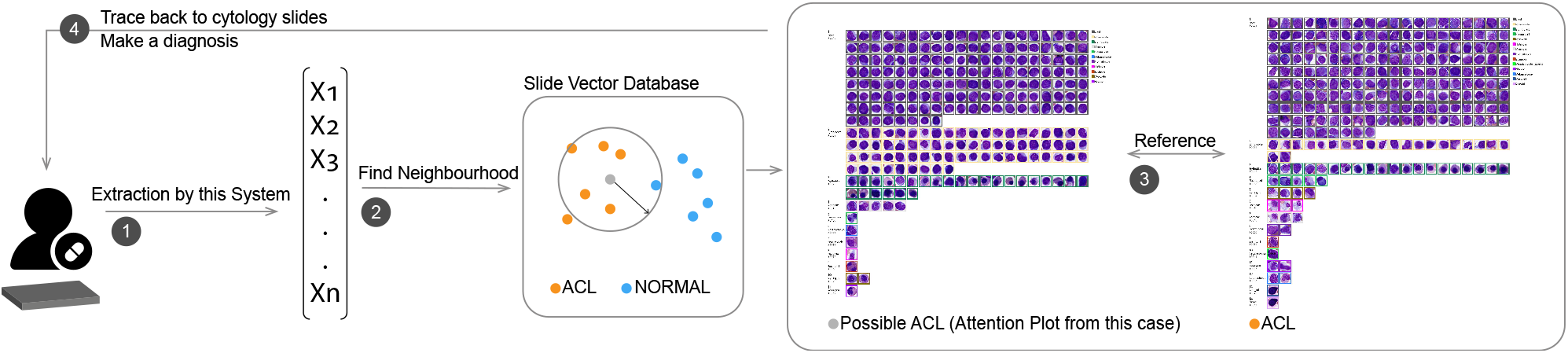
Proposed “AI as CDST” using this system. A patient’s cytology slide will be transformed as a slide vector by our system. Similar slides can be found in the database via vector search (Section 2.6) in a slide vector database. Then it may be predicted as ACL since its representation is inside the ACL “neighbourhood”. The attention plot of this slide can be used to see if it makes clinical sense. Furthermore, cells in the plot can be used as navigators that can trace back to the original WSI as the ultimate source. This way, the system can help doctors “digest” WSI information.

## 5 Data availability

The WSI that support the findings of this study are not publicly available due to them containing information that could compromise research participant privacy/consent. Source data underlying the main figures in the manuscript are available as Supplementary Data.

## 6 Code availability

Code for this system is available at https://zenodo.org/record/7150403

## 8 Author contributions

Youqing Mu designed and conducted experiments, analyzed data, created software and wrote the paper; H.R. Tizhoosh designed experiments, analyzed data, provided conceptual input, and contributed to writing the paper; Taher Dehkharghanian provided conceptual input and analyzed data; Clinton JV Campbell designed experiments, analyzed data, provided conceptual input and contributed to writing the paper.

## References

1. Pena, G. P. & Andrade-Filho, J. S. How does a pathologist make a diagnosis? Archives of pathology & laboratory medicine 133, 124–132 (2009).

2. Khoury, J. D. et al. The 5th edition of the World Health Organization classification of haematolymphoid tumours: myeloid and histiocytic/dendritic neoplasms. Leukemia, 1–17 (2022).

3. Alaggio, R. et al. The 5th edition of the World Health Organization classification of haematolymphoid tumours: lymphoid neoplasms. Leukemia, 1–29 (2022).

4. Rosai, J. The continuing role of morphology in the molecular age. Modern Pathology 14, 258–260 (2001).

5. Steiner, D. F., Chen, P.-H. C. & Mermel, C. H. Closing the translation gap: AI applications in digital pathology. Biochimica et Biophysica Acta (BBA)-Reviews on Cancer 1875, 188452 (2021).

6. Parwani, A. V. Next generation diagnostic pathology: use of digital pathology and artificial intelligence tools to augment a pathological diagnosis. Diagnostic Pathology 14, 1 (2019).

7. Williams, B. J., Bottoms, D. & Treanor, D. Future-proofing pathology: the case for clinical adoption of digital pathology. Journal of clinical pathology 70, 1010–1018 (2017).

8. Zarella, M. D. et al. A practical guide to whole slide imaging: a white paper from the digital pathology association. Archives of pathology & laboratory medicine 143, 222–234 (2019).

9. Pantanowitz, L. et al. Twenty years of digital pathology: an overview of the road travelled, what is on the horizon, and the emergence of vendor-neutral archives. Journal of pathology informatics 9, 40 (2018).

10. Sutton, R. T. et al. An overview of clinical decision support systems: benefits, risks, and strategies for success. NPJ digital medicine 3, 1–10 (2020).

11. Badillo, S. et al. An introduction to machine learning. Clinical pharmacology & therapeutics 107, 871–885 (2020).

12. Echle, A. et al. Deep learning in cancer pathology: a new generation of clinical biomarkers. British journal of cancer 124, 686–696 (2021).

13. Tran, K. A. et al. Deep learning in cancer diagnosis, prognosis and treatment selection. Genome Medicine 13, 1–17 (2021).

14. Jiang, Y., Yang, M., Wang, S., Li, X. & Sun, Y. Emerging role of deep learning-based artificial intelligence in tumor pathology. Cancer communications 40, 154–166 (2020).

15. Kumar, Y., Gupta, S., Singla, R. & Hu, Y.-C. A systematic review of artificial intelligence techniques in cancer prediction and diagnosis. Archives of Computational Methods in Engineering, 1–28 (2021).

16. Pudasaini, S. et al. Interpretation of bone marrow aspiration in hematological disorder. Journal of Pathology of Nepal 2, 309–312 (2012).

17. Tian, S. K. et al. Optimizing workflows and processing of cytologic samples for comprehensive analysis by next-generation sequencing: Memorial Sloan Kettering Cancer Center experience. Archives of pathology & laboratory medicine 140, 1200–1205 (2016).

18. Miller, D. C., Karcher, D. S., Kaul, K., et al. The crisis in the Pathology subspecialty fellowship application process: historical background and setting the stage. Academic pathology 9, 100030 (2022).

19. Sasada, K. et al. Inter-observer variance and the need for standardization in the morphological classification of myelodysplastic syndrome. Leukemia research 69, 54–59 (2018).

20. Font, P. et al. Inter-observer variance with the diagnosis of myelodysplastic syndromes (MDS) following the 2008 WHO classification. Annals of hematology 92, 19–24 (2013).

21. Naqvi, K. et al. Implications of discrepancy in morphologic diagnosis of myelodysplastic syndrome between referral and tertiary care centers. Blood, The Journal of the American Society of Hematology 118, 4690–4693 (2011).

22. Arber, D. A. et al. The 2016 revision to the World Health Organization classification of myeloid neoplasms and acute leukemia. Blood, The Journal of the American Society of Hematology 127, 2391–2405 (2016).

23. Tayebi, R. M. et al. Automated bone marrow cytology using deep learning to generate a histogram of cell types. Communications medicine 2, 1–14 (2022).

24. Ridgeway, J. A., Tinsley, S. & Kurtin, S. E. Practical guide to bone marrow sampling for suspected myelodysplastic syndromes. Journal of the Advanced Practitioner in Oncology 8, 29 (2017).

25. Font, P. et al. Interobserver variance in myelodysplastic syndromes with less than 5% bone marrow blasts: unilineage vs. multilineage dysplasia and reproducibility of the threshold of 2% blasts. Annals of hematology 94, 565–573 (2015).

26. Zhang, Z. et al. Pathologist-level interpretable whole-slide cancer diagnosis with deep learning. Nature Machine Intelligence 1, 236–245 (2019).

27. Audebert, N., Herold, C., Slimani, K. & Vidal, C. Multimodal deep networks for text and image-based document classification in Joint European Conference on Machine Learning and Knowledge Discovery in Databases (2019), 427–443.

28. Chen, Y., Gong, S. & Bazzani, L. Image search with text feedback by visiolinguistic attention learning in Proceedings of the IEEE/CVF Conference on Computer Vision and Pattern Recognition (2020), 3001–3011.

29. Van Tulder, G. & de Bruijne, M. Learning cross-modality representations from multi-modal images. IEEE transactions on medical imaging 38, 638–648 (2018).

30. Nagpal, K. et al. Development and validation of a deep learning algorithm for improving Gleason scoring of prostate cancer. NPJ digital medicine 2, 1–10 (2019).

31. Gorelick, L. et al. Prostate histopathology: Learning tissue component histograms for cancer detection and classification. IEEE transactions on medical imaging 32, 1804–1818 (2013).

32. Das, K., Conjeti, S., Roy, A. G., Chatterjee, J. & Sheet, D. Multiple instance learning of deep convolutional neural networks for breast histopathology whole slide classification in 2018 IEEE 15th International Symposium on Biomedical Imaging (ISBI 2018) (2018), 578–581.

33. Combalia, M. & Vilaplana, V. in Deep Learning in Medical Image Analysis and Multimodal Learning for Clinical Decision Support 274–281 (Springer, 2018).

34. Carbonneau, M.-A., Cheplygina, V., Granger, E. & Gagnon, G. Multiple instance learning: A survey of problem characteristics and applications. Pattern Recognition 77, 329–353 (2018).

35. Campanella, G. et al. Clinical-grade computational pathology using weakly supervised deep learning on whole slide images. Nature medicine 25, 1301–1309 (2019).

36. Kanavati, F. et al. Weakly-supervised learning for lung carcinoma classification using deep learning. Scientific reports 10, 1–11 (2020).

37. Tavolara, T. E. et al. Automatic discovery of clinically interpretable imaging biomarkers for Mycobacterium tuberculosis supersusceptibility using deep learning. EBioMedicine 62, 103094 (2020).

38. Tomita, N. et al. Attention-based deep neural networks for detection of cancerous and precancerous esophagus tissue on histopathological slides. JAMA network open 2, e1914645–e1914645 (2019).

39. Widrich, M. et al. Modern hopfield networks and attention for immune repertoire classification. Advances in Neural Information Processing Systems 33, 18832–18845 (2020).

40. Li, Z., Yuan, L., Xu, H., Cheng, R. & Wen, X. Deep multi-instance learning with induced self-attention for medical image classification in 2020 IEEE International Conference on Bioinformatics and Biomedicine (BIBM) (2020), 446–450.

41. Shao, Z. et al. Transmil: Transformer based correlated multiple instance learning for whole slide image classification. Advances in Neural Information Processing Systems 34, 2136–2147 (2021).

42. Niu, Z., Zhong, G. & Yu, H. A review on the attention mechanism of deep learning. Neurocomputing 452, 48–62 (2021).

43. Lu, M. Y. et al. Data-efficient and weakly supervised computational pathology on whole-slide images. Nature biomedical engineering 5, 555–570 (2021).

44. Raskin, R. E. & Messick, J. B. Bone marrow cytologic and histologic biopsies: indications, technique, and evaluation. Veterinary Clinics: Small Animal Practice 42, 23–42 (2012).

45. Gilotra, M., Gupta, M., Singh, S. & Sen, R. Comparison of bone marrow aspiration cytology with bone marrow trephine biopsy histopathology: An observational study. Journal of Laboratory Physicians 9, 182–189 (2017).

46. Mu, Y. et al. A BERT model generates diagnostically relevant semantic embeddings from pathology synopses with active learning. Communications medicine 1, 1–13 (2021).

47. Parsons, V. L. Stratified sampling. Wiley StatsRef: Statistics Reference Online, 1–11 (2014).

48. Bochkovskiy, A., Wang, C.-Y. & Liao, H.-Y. M. YOLOv4: Optimal Speed and Accuracy of Object Detection. arXiv e-prints, arXiv–2004 (2020).

49. Duan, K. et al. CenterNet: Keypoint Triplets for Object Detection. arXiv e-prints, arXiv–1904 (2019).

50. Hopfield, J. J. Neural networks and physical systems with emergent collective computational abilities. Proceedings of the national academy of sciences 79, 2554–2558 (1982).

51. Ramsauer, H. et al. Hopfield networks is all you need. arXiv preprint arXiv:2008.02217 (2020).

52. Vaswani, A. et al. Attention is all you need. Advances in neural information processing systems 30 (2017).

53. Musgrave, K., Belongie, S. & Lim, S.-N. PyTorch Metric Learning 2020. arXiv: 2008.09164 [cs.CV].

54. Yang, L. & Jin, R. Distance metric learning: A comprehensive survey. Michigan State Universiy 2, 4 (2006).

55. Schroff, F., Kalenichenko, D. & Philbin, J. Facenet: A unified embedding for face recognition and clustering in Proceedings of the IEEE conference on computer vision and pattern recognition (2015), 815–823.

56. Hoffer, E. & Ailon, N. Deep metric learning using triplet network in International workshop on similarity-based pattern recognition (2015), 84–92.

57. Sohn, K. Improved deep metric learning with multi-class n-pair loss objective. Advances in neural information processing systems 29 (2016).

58. Chen, B. & Deng, W. Deep embedding learning with adaptive large margin N-pair loss for image retrieval and clustering. Pattern Recognition 93, 353–364 (2019).

59. Khosla, P. et al. Supervised contrastive learning. Advances in Neural Information Processing Systems 33, 18661–18673 (2020).

60. De Boer, P.-T., Kroese, D. P., Mannor, S. & Rubinstein, R. Y. A tutorial on the cross-entropy method. Annals of operations research 134, 19–67 (2005).

61. Platzer, C. & Dustdar, S. A vector space search engine for web services in Third European Conference on Web Services (ECOWS’05) (2005), 9–pp.

62. Musgrave, K., Belongie, S. & Lim, S.-N. A metric learning reality check in European Conference on Computer Vision (2020), 681–699.

63. Fix, E. & Hodges, J. L. Discriminatory analysis. Nonparametric discrimination: Consistency properties. International Statistical Review/Revue Internationale de Statistique 57, 238–247 (1989).

64. Pedregosa, F. et al. Scikit-learn: Machine Learning in Python. Journal of Machine Learning Research 12, 2825–2830 (2011).

65. Xu, Q.-S. & Liang, Y.-Z. Monte Carlo cross validation. Chemometrics and Intelligent Laboratory Systems 56, 1–11 (2001).

66. Abdulrahman, A. A. et al. Is a 500-cell count necessary for bone marrow differentials? A proposed analytical method for validating a lower cutoff. American journal of clinical pathology 150, 84–91 (2018).

67. Lee, S.-H. et al. ICSH guidelines for the standardization of bone marrow specimens and reports. International journal of laboratory hematology 30, 349–364 (2008).

68. Bain, B. J. & Ahmad, S. Chronic neutrophilic leukaemia and plasma cell-related neutrophilic leukaemoid reactions. British Journal of Haematology 171, 400–410 (2015).

69. Srivastava, N., Hinton, G., Krizhevsky, A., Sutskever, I. & Salakhutdinov, R. Dropout: a simple way to prevent neural networks from overfitting. The journal of machine learning research 15, 1929–1958 (2014).

70. Wang, J., Perez, L., et al. The effectiveness of data augmentation in image classification using deep learning. Convolutional Neural Networks Vis. Recognit 11, 1–8 (2017).

71. Hestness, J. et al. Deep Learning Scaling is Predictable, Empirically. arXiv e-prints, arXiv–1712 (2017).

72. Xu, W. Toward human-centered AI: a perspective from human-computer interaction. interactions 26, 42–46 (2019).

73. Li, M. et al. Clip-event: Connecting text and images with event structures in Proceedings of the IEEE/CVF Conference on Computer Vision and Pattern Recognition (2022), 16420–16429.

