## Supplement Data for "Whole slide image representation in bone marrow cytology"

| Experiment | Hopfield on Cell Bags | rHCT | AvgPooling on Cell Bags | Random |
|---|---|---|---|---|
| 0 | 0.606 | 0.545 | 0.508 | 0.382 |
| 1 | 0.593 | 0.545 | 0.511 | 0.375 |
| 2 | 0.582 | 0.551 | 0.499 | 0.372 |
| 3 | 0.545 | 0.546 | 0.492 | 0.402 |
| 4 | 0.589 | 0.566 | 0.529 | 0.395 |

Table S1: The mAP@5 comparison.

Figure S1: Whole slide images' embeddings from average pooling. 2D projection of WSI embeddings from the 307 cases in the test set is shown. Comparing to the results from Hopfield pooling, its WSI embeddings form fewer distinct clusters.

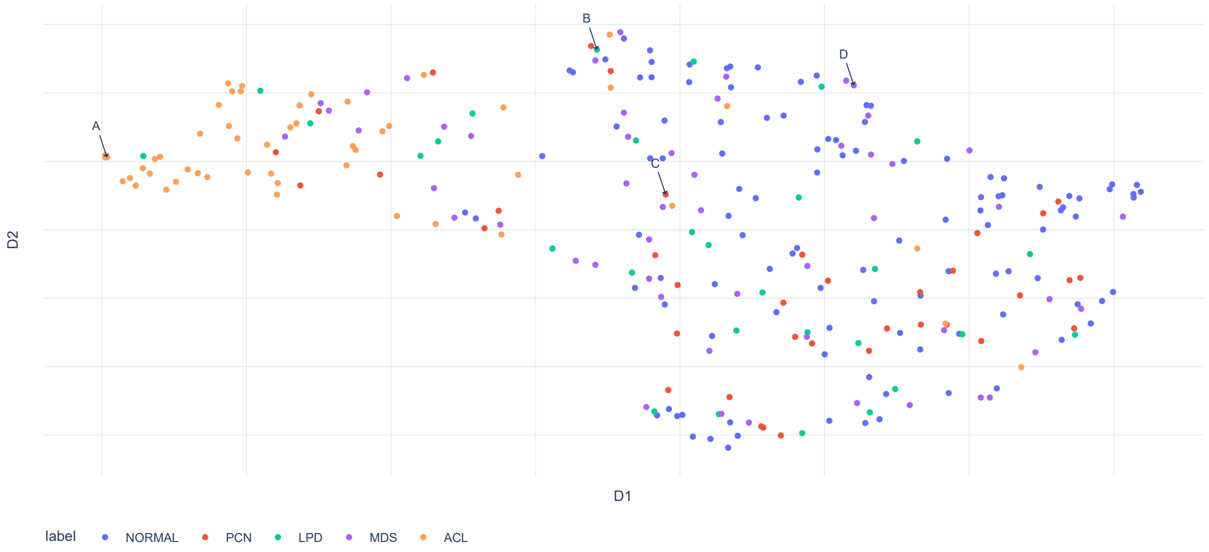

| Label | F1 Score | Precision | Recall | Method |
|---|---|---|---|---|
| ACL | 0.711±0.039 | 0.693±0.082 | 0.737±0.030 | AvgPooling on Cell Bags |
| LPD | 0.039±0.058 | 0.300±0.447 | 0.021±0.031 | AvgPooling on Cell Bags |
| MDS | 0.126±0.069 | 0.290±0.140 | 0.081±0.046 | AvgPooling on Cell Bags |
| NORMAL | 0.652±0.014 | 0.505±0.004 | 0.919±0.045 | AvgPooling on Cell Bags |
| PCN | 0.067±0.033 | 0.172±0.066 | 0.043±0.024 | AvgPooling on Cell Bags |
| ACL | 0.196±0.063 | 0.192±0.062 | 0.200±0.064 | Empirical Inference |
| LPD | 0.123±0.022 | 0.111±0.020 | 0.138±0.024 | Empirical Inference |
| MDS | 0.189±0.026 | 0.227±0.031 | 0.162±0.022 | Empirical Inference |
| NORMAL | 0.418±0.053 | 0.403±0.051 | 0.433±0.054 | Empirical Inference |
| PCN | 0.083±0.056 | 0.086±0.057 | 0.081±0.054 | Empirical Inference |
| ACL | 0.788±0.015 | 0.804±0.019 | 0.773±0.022 | rHCT |
| LPD | 0.285±0.045 | 0.577±0.098 | 0.193±0.046 | rHCT |
| MDS | 0.315±0.067 | 0.566±0.089 | 0.219±0.052 | rHCT |
| NORMAL | 0.675±0.011 | 0.527±0.009 | 0.936±0.015 | rHCT |
| PCN | 0.176±0.097 | 0.666±0.258 | 0.103±0.059 | rHCT |
| ACL | 0.737±0.050 | 0.742±0.083 | 0.737±0.055 | Hopfield on Cell Bags |
| LPD | 0.419±0.092 | 0.563±0.069 | 0.338±0.093 | Hopfield on Cell Bags |
| MDS | 0.331±0.057 | 0.372±0.046 | 0.300±0.066 | Hopfield on Cell Bags |
| NORMAL | 0.666±0.017 | 0.585±0.024 | 0.776±0.046 | Hopfield on Cell Bags |
| PCN | 0.481±0.047 | 0.643±0.114 | 0.389±0.041 | Hopfield on Cell Bags |

Table S2: The performance comparison between full system, average pooling and empirical inference. We also show the standard deviation computed across five experiments.
